# Metformin Improves Ovarian Function and Increases Egg Production in Broiler Breeder Hens

**DOI:** 10.1101/2022.07.13.499919

**Authors:** Evelyn A. Weaver, Ramesh Ramachandran

## Abstract

Broiler breeder hens, the parent stock of commercial broiler chickens, have poor reproductive efficiency associated with aberrant and excessive recruitment of ovarian follicles which results in sub-optimal egg production, fertility, and hatchability. The reproductive dysfunction observed in these hens resembles polycystic ovary syndrome in women, a condition wherein metformin is prescribed as a treatment. The main objectives of this study were to determine the effect of metformin on body weight, abdominal fat pad weight, ovarian function, and plasma steroid hormone concentrations. Broiler breeder hens were treated with 0, 25, 50, or 75 mg/kg body weight of metformin mixed in the diet for 40 weeks (n= 45 hens/treatment; 25-65 weeks of age). At 65 weeks of age, hens that received the highest dose of metformin had significantly lower body and abdominal fat pad weights (p < 0.05) than the control. Metformin treatment, at all levels, normalized the preovulatory and prehierarchical ovarian follicular hierarchy. Metformin (50 or 75 mg/kg body weight) significantly increased the total number of eggs laid per hen during the entire production period and these hens had significantly greater fertility and hatchability at 65 weeks of age compared to the control (p < 0.05). Metformin treatment at all levels altered the plasma profile of reproductive hormones, with significantly lower plasma testosterone concentrations and a decreased testosterone to androstenedione ratio in hens that received metformin (p < 0.05). Future studies should focus on the mechanisms underlying the beneficial effects of metformin in improving the reproductive efficiency of broiler breeder hens.

**In Brief:** The pathophysiology of the ovarian dysfunction encountered in broiler breeder hens remains poorly understood but is similar to a condition in women known as PCOS. This study reveals that metformin may provide a cheap and effective method of improving ovarian function in broiler breeder hens.

## INTRODUCTION

Broiler breeder hens, the parent stock of commercial broiler chickens, have poor reproductive efficiency due to the selection pressure for superior growth-related traits in broiler progeny. Decades of intensive genetic selection for growth-related traits negatively associated with reproductive function have led to severe ovarian dysfunction in the broiler breeder hen (De Jong and Guémené, 2011; Zuidhof et al., 2014). Full-fed broiler breeder hens accumulate excessive amounts of visceral adipose tissue and often have heavier ovaries, excessive follicular recruitment, double ovulation, or increased incidence of ovarian regression (Yu et al., 1992). Consequently, broiler breeder hens have decreased egg production, sub-optimal fertility and hatchability, and decreased viability of embryos (Yu et al., 1992).

The pathophysiology underlying the ovarian dysfunction in broiler breeder hens is still unknown, but previous studies have shown that it may be related to obesity-induced metabolic dysfunction and lipotoxicity (Cassy et al., 2004; Chen et al., 2006; Pan et al., 2012; Walzem and Chen, 2014). In studies where broiler breeder hens were allowed to consume feed *ad libitum*, significantly higher body weights, increased abdominal fat pad weights, and increased liver weights were observed (Chen et al., 2006; Taherkhani et al., 2010; Pan et al., 2012). Such increases in the accumulation of abdominal adipose tissue and increased weight gain were associated with increased plasma glucose, insulin, non-esterified fatty acids, triacylglycerol, cholesterol, phospholipid, and ceramide levels (Chen et al., 2006; Taherkhani et al., 2010; Pan et al., 2012). An increased incidence of ovarian dysfunction, follicle atresia, early ovarian regression, and altered plasma progesterone and estradiol levels was also observed in these hens (Cassy et al., 2004; Chen et al., 2006; Taherkhani et al., 2010; Pan et al., 2012). In a study where broiler breeder hens were treated with testosterone, higher rates of internal ovulation and increased disruption of the reproductive cycle was observed (Navara, Kristen J et al., 2015).

These findings suggest that increased adiposity and disrupted steroid hormone signaling may play a central role in developing ovarian dysfunction. Although previous studies have focused on the effects of *ad libitum* feeding, the hyper-recruitment of prehierarchical ovarian follicles and a deranged preovulatory follicular hierarchy are frequently encountered even when the hens are fed a restricted amount of feed.

Interestingly, the abnormal ovarian changes and disturbed plasma profiles of reproductive hormones observed in broiler breeder hens appear similar to a condition in women known as polycystic ovary syndrome (PCOS). PCOS is most commonly associated with obesity, metabolic syndrome, irregular steroid hormone production, irregular ovulation, and poor fertility, all of which have been previously observed in broiler breeder hens (Pasquali et al., 2003; Walzem and Chen, 2014). Metformin, a biguanide compound, is the most commonly prescribed drug to treat polycystic ovary syndrome. In women with PCOS, metformin has been shown to reduce insulin resistance and correct metabolic disturbances commonly observed in PCOS. Metformin has also been associated with increased ovulation, fertilization, and pregnancy rates (Velazquez et al., 1997; Fleming et al., 2002; Nestler et al., 2002; Tang et al., 2006b; Morley et al., 2017; Tso et al., 2020), normalization of the endocrine profile, as well as a return to normal menstrual cyclicity (Velazquez et al., 1994; Morin-Papunen et al., 1998; Moghetti et al., 2000; van Santbrink et al., 2005; Xing et al., 2020). While this improvement in reproductive function has previously been associated with the correction of insulin resistance (Perriello et al., 1994; Prager et al., 1986; Jakubowicz et al., 2001) and improved energy metabolism, other studies have shown that metformin can exert direct effects on the ovary (Ehrmann et al., 1997; Vrbikova et al., 2001; Bertoldo et al., 2014). In agreement with previous studies, we found that metformin attenuates steroidogenesis in the granulosa cells isolated from broiler breeder hen ovarian follicles (Weaver and Ramachandran, 2020).

Based on the preceding, we hypothesized that dietary metformin supplementation would improve reproductive efficiency in the broiler breeder hen. The main objectives of the present study are to determine the effects of metformin on (i) body weight and abdominal fat pad weight, (ii) ovarian prehierarchical and preovulatory follicular hierarchy, (iii) egg production, fertility, and hatchability, and (iv) plasma concentrations of steroid hormones.

## MATERIALS AND METHODS

### Animal Husbandry and Metformin Treatment

The Pennsylvania State University’s Institutional Animal Care and Use Committee approved all animal procedures described herein (protocol number PRAMS200746656). The overall experimental design and sample collection timeline are provided in Supplementary Figure 1. Broiler breeder female (Cobb 500; Cobb-Vantress Inc.) and male (Hubbard M99) chickens were maintained from 1 day-old until the completion of the study at the Poultry Education and Research Center at The Pennsylvania State University (University Park, PA). Broiler breeder hens and roosters were housed individually in battery cages from 21 weeks of age and were feed-restricted according to the Cobb Breeder and Hubbard management guide. Feed restriction began at 2 weeks of age and continued through the end of the study (65 weeks of age). While rearing, all birds were weighed weekly to adjust feed amounts and to maintain uniformity. A set feed amount of 150 g/hen/day was fed from 25-65 weeks of age. Water was provided *ad libitum*. The chickens were photo stimulated with a 16h light:8h dark (4:00 to 20:00 hours) cycle for the study duration. At 25 weeks of age, the broiler breeder female chickens were randomly allocated to four experimental groups (n=45) to receive four metformin levels (0, 25, 50, or 75 mg/kg body weight; Midwest Veterinary Supply, Lakeville, MN) supplemented in the diet. The metformin treatment continued until the end of the study, at which the chickens were 65 weeks of age. The doses of metformin were chosen based on preliminary studies conducted in our laboratory. Broiler breeder hens were supplemented with various doses of metformin (25-200 mg metformin/kg body weight). We found that supplementing ≥ 100 mg metformin/kg body weight resulted in negative outcomes with most of the hens losing a substantial amount of body weight and fat pad leading many to cease laying. Thus, we chose to utilize metformin doses less than 100 mg metformin/kg body weight. Male chickens received no metformin supplementation in their diets. The broiler breeder hens were on a time-restricted feeding schedule, where a fixed amount of feed was given each morning as recommended by Cobb-Vantress Inc. This fixed amount of feed and once-a-day feeding schedule ensured that each hen consumed the correct dose of metformin daily. A random subset of broiler breeder hens from each treatment group (n=10) was weighed every ten weeks to adjust the amount of metformin mixed into the feed according to their change in weights over time. Egg production was recorded daily for each broiler breeder hen starting at 22 weeks of age through the end of the study.

### Body weight

A random subset of hens (n=10/treatment group) were weighed every ten weeks starting at 30 weeks of age for the duration of the study to determine the effect of metformin on body weight. The body weights were also used to adjust the amount of metformin added to the feed to maintain the doses at 0, 25, 50, or 75 mg/kg throughout the study. A subset of hens (n=12 hens/treatment group) were weighed at the end of the study (65 weeks of age). At 65 weeks, broiler breeder hens were euthanized and the abdominal fat pad was collected and weighed (n= 12 hens/treatment group).

### Fertility and hatchability

For fertility and hatchability studies, a subset of female chickens from each experimental group was randomly selected (n=20/treatment group). The same chickens were utilized in all fertility and hatchability data collections. For artificial insemination (AI), semen was collected and pooled from 15 broiler breeder male chickens. Immediately following collection, 50 mL of undiluted, pooled semen was deposited into the oviduct of each broiler breeder female chicken. Inseminations were done every other day, and eggs were collected daily and stored at 55°F until the end of a 10-day collection period (n= 100-150 eggs/treatment group). Fertility and hatchability data collections were conducted at seven different time points throughout the study (35, 40, 45, 50, 55, 60, and 65 weeks of age). Eggs were placed in Chick Master (Medina, OH) incubators set at 100°F and 58-60% humidity. Eggs were candled on day 10 of incubation to determine fertility for each individual hen (n= 20 hens/treatment group), represented as the total number of fertile eggs to the total number of eggs set for each hen. On day 19 of incubation, fertile eggs were transferred to Chick Master hatchers set at 98.2°F with humidity ~65-70%. Hatchability was determined on day 21 and is represented as the total number of chicks hatched to the total number of fertile eggs transferred to the hatchers for each individual hen.

### Plasma Steroid Hormone Quantification

Blood samples were collected from the wing vein from a subset of fasted hens (n=6/ treatment group) every five weeks to quantify plasma steroid hormone concentrations. Blood samples were immediately placed on ice and centrifuged within 1 h at 2300 x g for 15 min at 4°C. Plasma was collected and stored at −80°C for quantification of progesterone, total estrogens, testosterone, and androstenedione concentrations. Plasma progesterone concentration was determined by a commercial human double-antibody radioimmunoassay (RIA) kit (cat# 0717010, MP Biomedicals, LLC, Irvine, CA) according to the manufacturer’s recommendations.

The intraassay coefficient of variation (CV) was 8%, and an interassay CV of 9% with a sensitivity of 0.2-50 ng/mL for this assay. For the analysis of plasma total estrogens, testosterone, and androstenedione, an extraction step was necessary prior to running the assay according to the manufacturer’s recommendations. Briefly, 6 mL of ethyl acetate: hexane, mixed in a ratio of 3:2, was added to 500 mL of plasma. Tubes were vortexed vigorously for 60 seconds, and phases were allowed to separate. Next, 5 ml of the organic phase (top phase) was withdrawn and evaporated under nitrogen gas. The sample residue was reconstituted in 2.5 mL of steroid diluent provided by the manufacturer and incubated at room temperature for one hour while gently mixed on a plate shaker. Extracted samples were stored at −20°C. According to the manufacturer’s recommendations, the total estrogens plasma concentration was determined by a commercial human double-antibody RIA kit (cat# 07140205, MP Biomedicals). For the total estrogens assay, the intraassay and interassay CV were 8% and 7%, respectively, while the sensitivity of this assay was 2.5-100 pg/mL. Plasma testosterone and androstenedione concentrations were determined by RIA using commercial reagents (cat# 0718910 and cat# 07109202, MP Biomedicals, LLC, Irvine, CA). For testosterone and androstenedione, intraassay CVs were 8%, and the interassay CVs were 3% and 12%, respectively. The sensitivity of the testosterone and androstenedione assays were 0.025-10 ng/mL and 0.1-10 ng/mL, respectively.

The commercial radioimmunoassay kits were previously validated and used in our laboratory (Ocón-Grove et al., 2007) and proper controls were utilized in each run.

### Statistical Analyses

All data were subjected to a one-way analysis of variance (ANOVA) using GraphPad Prism version 9.1.0, GraphPad Software, San Diego, CA. Differences among individual means were determined by using Tukey’s HSD multiple comparisons test. To analyze cumulative egg production and fertility and hatchability per hen between the treatment groups and over time, data were subjected to a repeated measures two-way ANOVA using GraphPad Prism version 9.1.0, (GraphPad Software, San Diego, CA). Differences among individual means were determined by using Tukey’s HSD multiple comparisons test. A chi-square test was performed to determine the statistical significance of the number of hens still laying at the end of the trial (metformin treatment vs. control). A probability level of P < 0.05 is considered statistically significant. All data are represented as mean ± standard error of the mean. All data points are compared within a timepoint and not across timepoints, except for the radioimmunoassay data, which was also analyzed across time for each treatment group.

## RESULTS

### Metformin decreases the body weight and accretion of abdominal adipose tissue

Body weight of all hens at 20- or 30 weeks of age did not differ. Hens receiving 75 mg metformin/kg body weight had significantly lower body weights at 40 weeks of age when compared with hens receiving 25 mg metformin/kg body weight. At 50 and 60 weeks of age, hens receiving the highest dose of metformin had significantly lower body weights when compared to all other treatment groups. However, hens receiving the highest dose only differed from those receiving 0- or 25 mg/kg metformin/kg body weight at 65 weeks of age (Fig. 1A).

**Figure 1.**
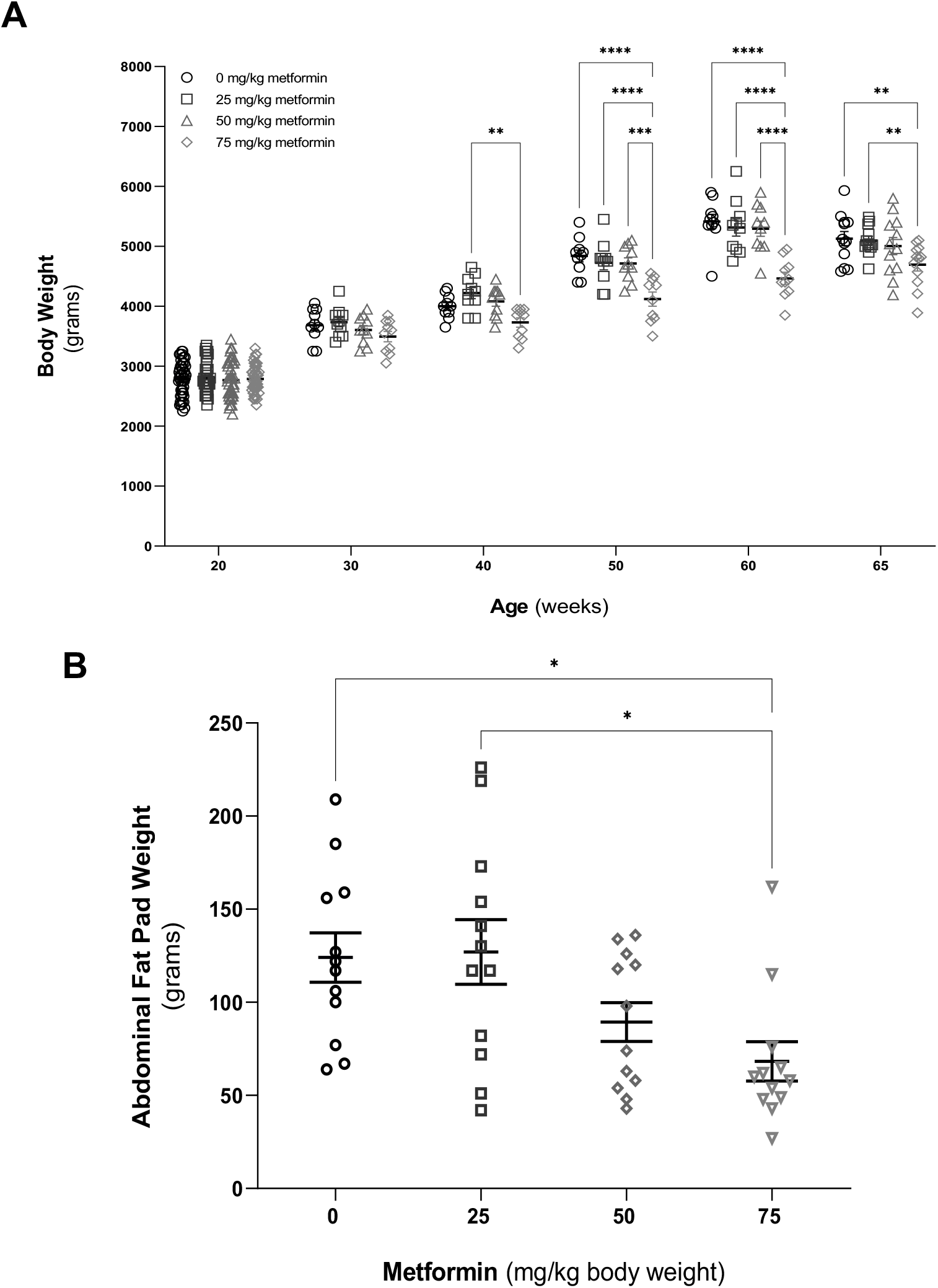
Effect of metformin supplementation on body weight (A) and abdominal fat pad weight (B). Broiler breeder hen diet was supplemented with metformin (0, 25, 50, or 75 mg/kg bw) from 25 to 65 weeks of age. All hens (n=45 hen/treatment group) were weighed at 25 weeks of age, and a random subset (n=10 hens/treatment group) were weighed every 10 weeks from 30 to 65 weeks of age. The entire abdominal fat pad was collected during necropsy done at the end of study. Values are means ± SEM. * *P* ≤ 0.05, ** *P* ≤ 0.01, *** *P* ≤ 0.001, **** *P* ≤ 0.0001 by one-way ANOVA.

Consistent with the observed decrease in body weight, hens receiving the highest dose of metformin had significantly lower abdominal fat pad weights at the end of the study (65 weeks of age) compared to the control group and the group receiving the lowest dose of metformin (25 mg/kg of body weight) (Fig. 1B).

### Metformin affects the ovarian follicular hierarchy, increases overall egg production and maintains fertility and hatchability

Figures 2A and 2B show representative a hierarchy of preovulatory and prehierarchical ovarian follicles from a hen (60 weeks of age) that received 0 or 25 mg/kg body weight of metformin, respectively. As seen in Fig. 2A, there are an excessive number of preovulatory and prehierarchical follicles in the hens that received no metformin. In contrast, supplementation of metformin in the diet restores the ovarian follicular hierarchy towards a normal phenotype (Fig. 2B). In addition to the changes in the follicular hierarchy, hens which received 50 or 75 mg/kg metformin laid a significantly greater number of eggs per hen on average when compared to the control, starting at 50 weeks of age through the end of the study (Fig 3A). When cumulative egg production per hen was observed over the entire production period (22-65 weeks of age), hens that received 50 or 75 mg metformin /kg body weight laid a significantly greater number of eggs per hen when compared to the control (Fig. 3B), which is reflected in a greater total number of eggs laid per treatment group over the entire study (Fig. 3B insert). Furthermore, hens that received the highest dose of metformin had a significantly greater number of hens remaining in production at the end of the study when compared to the control (Table 1). Interestingly, hens which received 50 or 75 mg metformin/kg body weight had significantly greater fertility (Fig. 4A) and hatchability (Fig. 4B) percentages at 65 weeks of age when compared to hens which received no metformin in the diet.

**Figure 2.**
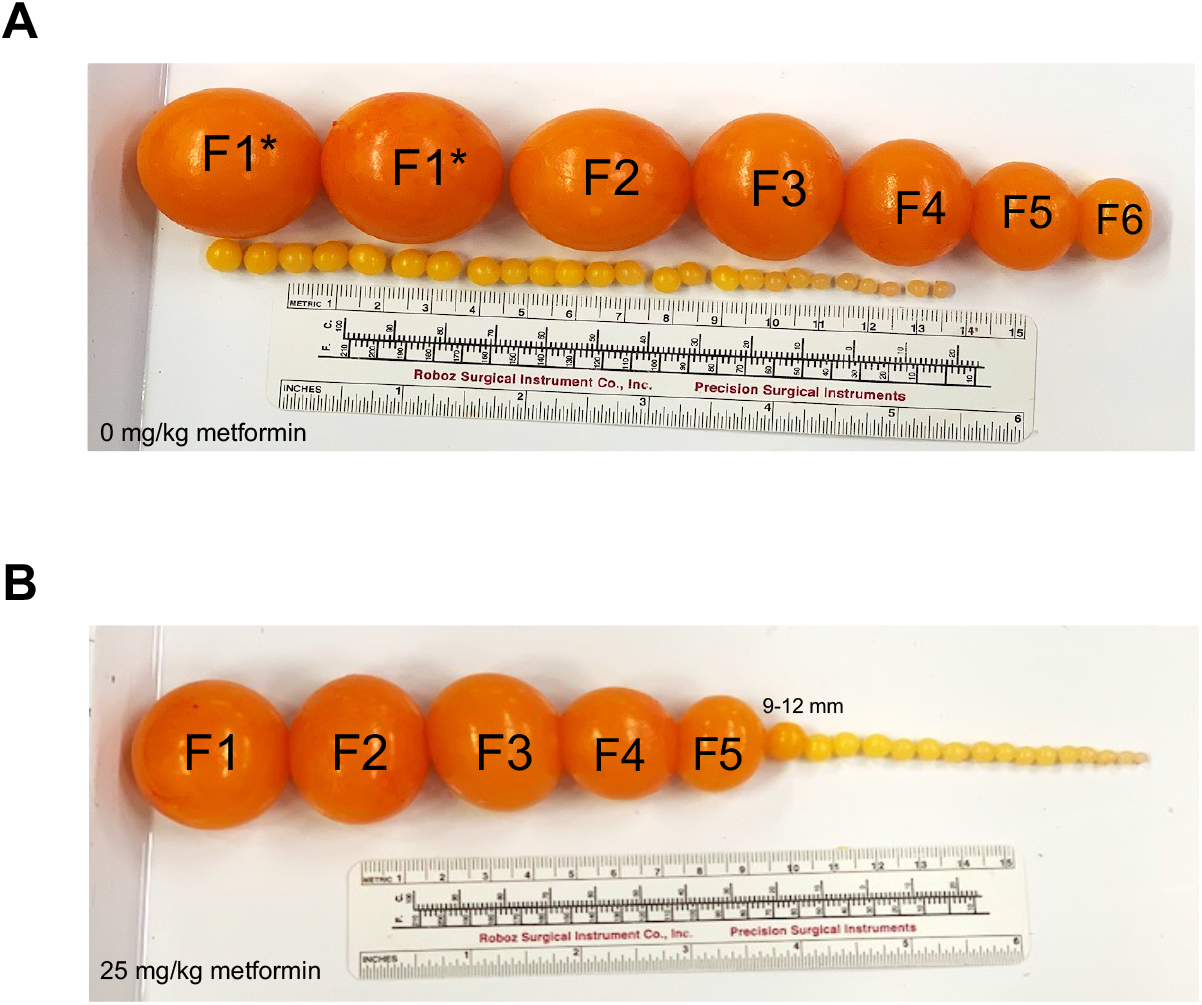
Effect of metformin supplementation on the ovarian follicular hierarchy. Broiler breeder hen diet was supplemented with metformin (0, 25, 50, or 75 mg/kg bw) from 25 to 65 weeks of age. Representative photographs of the ovarian follicular hierarchy from broiler breeder hens (60 weeks of age) that received 0 or 25 mg/ kg body weigh metformin (A and B, respectively). A double hierarchy of preovulatory follicles (denoted F1-F5) and hyper-recruitment of 3-5 mm and 6-8 mm sized prehierarchical follicles are seen in **(A)** while hens which received 25 mg metformin/kg body weight display a normal hierarchy of preovulatory and prehierarchical follicles in **(B)**.

**Figure 3.**
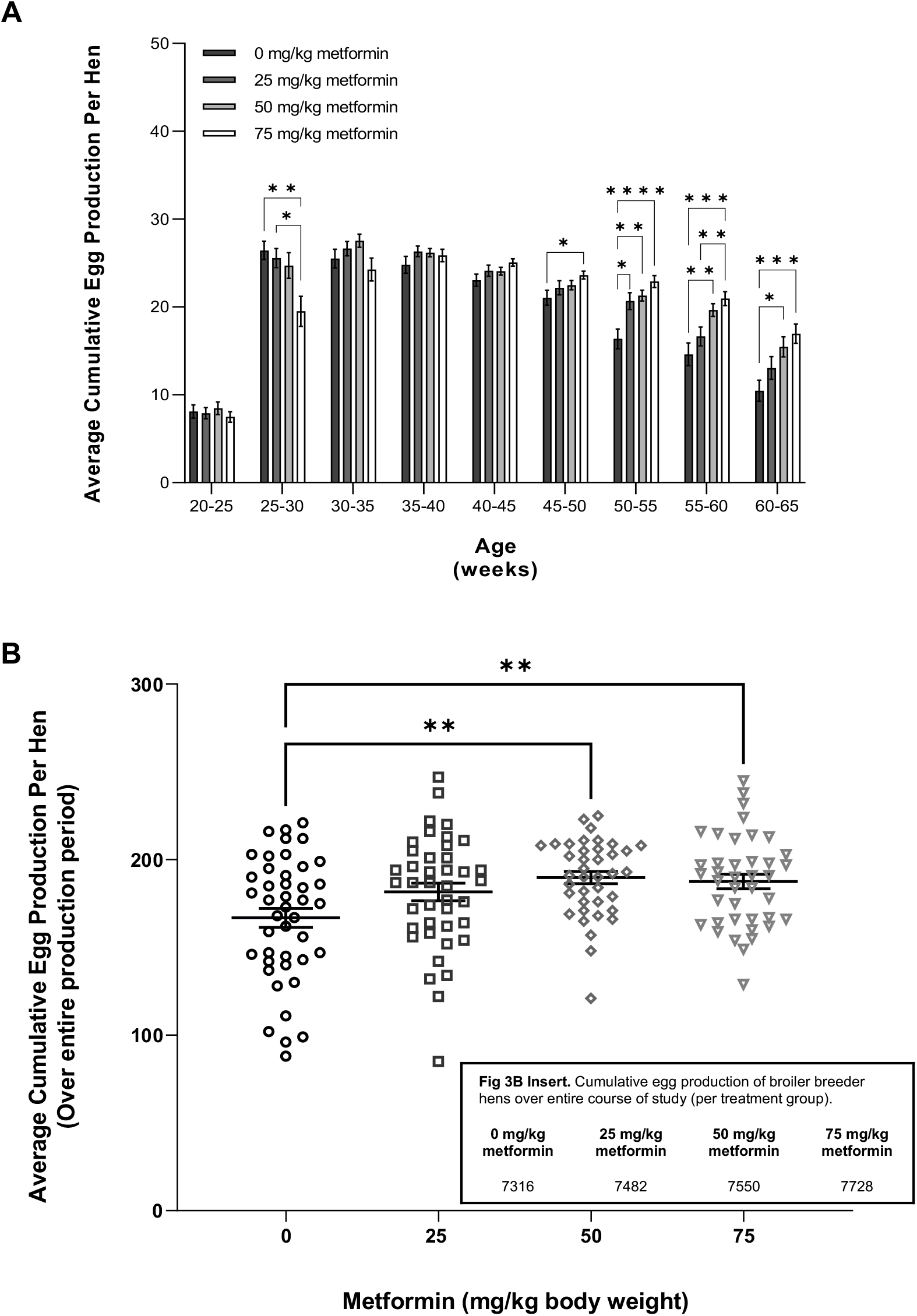
Effect of metformin supplementation on egg production. Broiler breeder hen diet was supplemented with metformin (0, 25, 50, or 75 mg/kg bw) from 25 to 65 weeks of age. Egg production was recorded daily for each hen from 22 weeks of age through 65 weeks of age (n=40-45 hens/treatment group). Average cumulative egg production per hen every 5 weeks over entire course of the study **(A)** and the overall average cumulative egg production per hen **(B).** Values are means ± SEM. * *P* ≤ 0.05, ** *P* ≤ 0.01, *** *P* ≤ 0.001, **** *P* ≤ 0.0001 by repeated measures two-way ANOVA.

**Figure 4.**
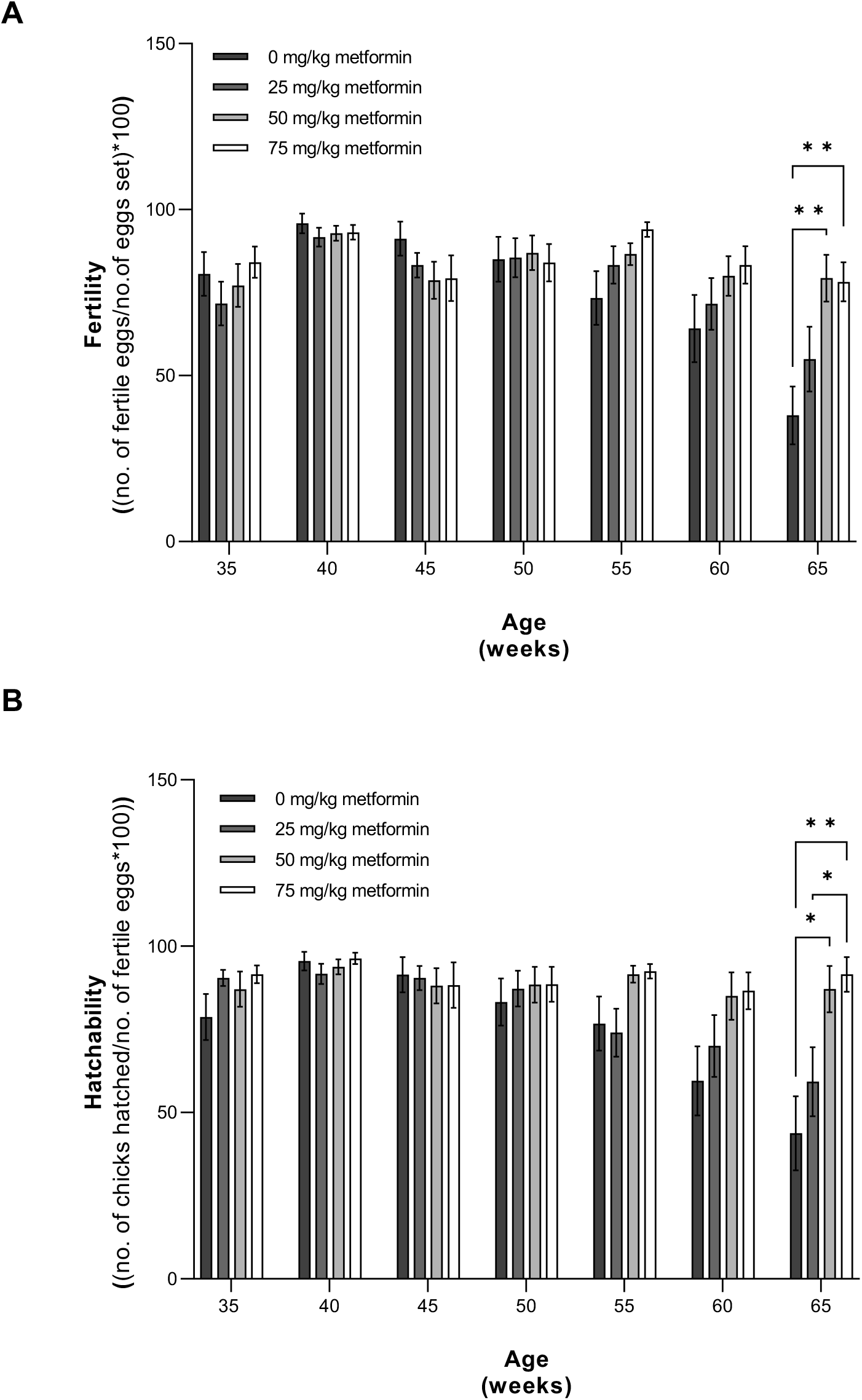
Effect of metformin supplementation on fertility and hatchability. Broiler breeder hen diet was supplemented with metformin (0, 25, 50, or 75 mg/kg bw) from 25 to 65 weeks of age. For fertility and hatchability studies, a subset of female chickens from each experimental group (n=20/treatment group) were artificially inseminated using pooled semen. Eggs were candled on day 10 of incubation to determine fertility for each individual hen (n= 20 hens/treatment group), represented as the total number of fertile eggs to the total number of eggs set for each hen. On day 19 of incubation, fertile eggs were transferred to Chick Master hatchers. Hatchability was determined on day 21 and is represented as the total number of chicks hatched to the total number of fertile eggs transferred to the hatchers for each individual hen. Average fertility **(A)** and average hatchability **(B).** Values are means ± SEM. * *P* ≤ 0.05, ** *P* ≤ 0.01 by repeated measures two-way ANOVA.

**Figure 5.**
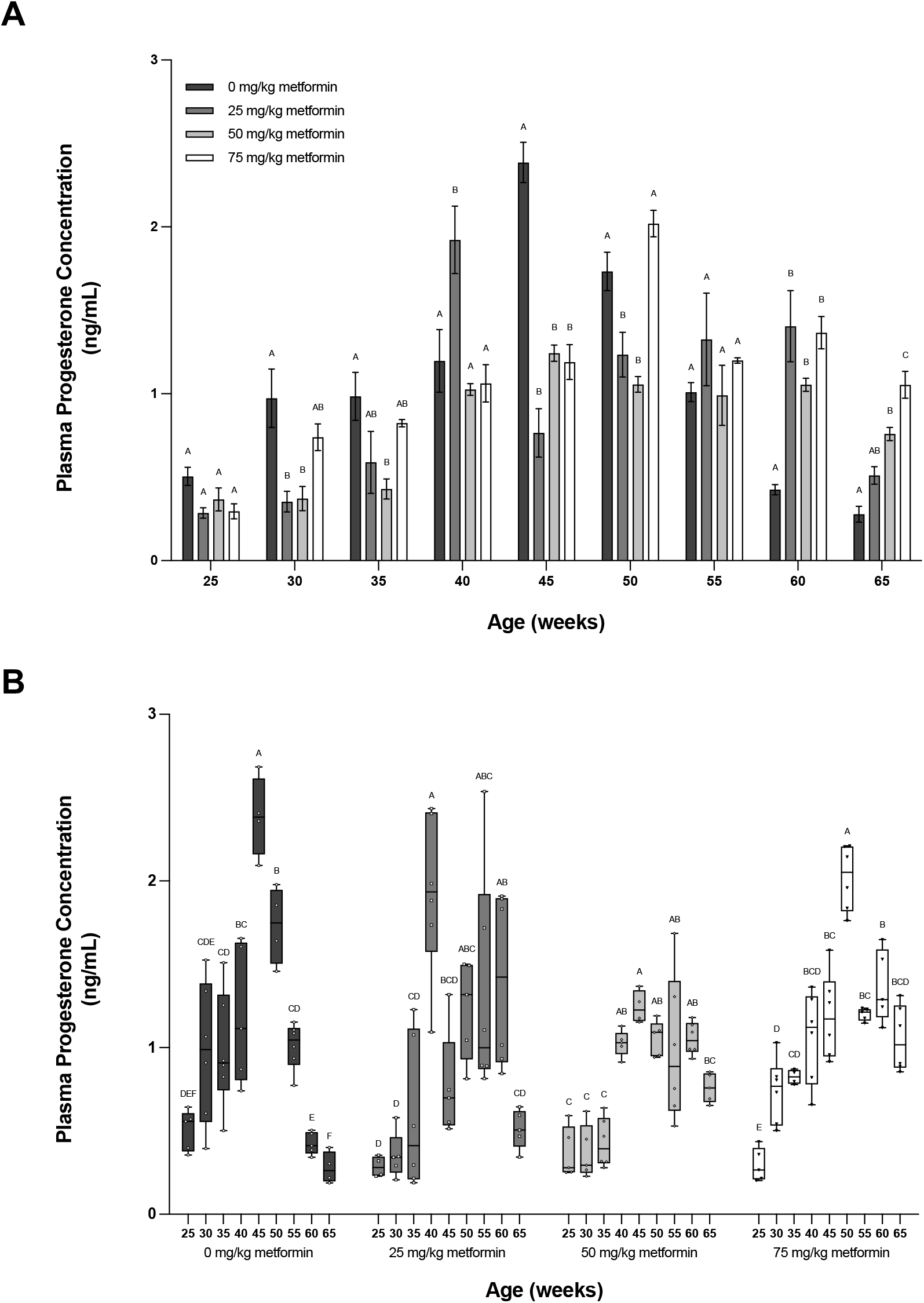
Effect of metformin supplementation on plasma progesterone concentration. Broiler breeder hen diet was supplemented with metformin (0, 25, 50, or 75 mg/kg bw) from 25 to 65 weeks of age. Progesterone concentrations (ng/mL) were determined by radioimmunoassay in blood samples collected from a subset of hens (n=6 hens/treatment group) every 5 weeks from 25-65 weeks of age **(A)**. Plasma progesterone concentrations (ng/ml) within each treatment group over the course of the study is shown in **(B)**. Values are means ± SEM. ^A, B, C, D, E, F^ P < 0.05 by one-way ANOVA; n= 6 hens/treatment group for each time point.

**Table 1:** Effect of metformin on number of broiler breeder hens still laying at end of trial. Egg production was recorded daily for each individual hen beginning at 22 weeks of age through 65 weeks of age (n= 43-45 hens). Hens were considered to still be in lay if they laid ≥ 3 eggs/week.

|  | 0 mg/kg<br>metformin | 25 mg/kg<br>metformin | 50 mg/kg<br>metformin | 75 mg/kg<br>metformin |
| --- | --- | --- | --- | --- |
| **% of Hens**<br>**Still**<br>**Laying**<br>(end of trial) | 44.44%<sup>A</sup><br>(20/45) | 59.09%<sup>AB</sup><br>(26/44) | 63.64%<sup>AB</sup><br>(28/44) | 69.77%<sup>B</sup><br>(30/43) |

<sup>A,B</sup> Values with different letters are significantly different by Chi-square statistical test ( $P \leq 0.05$ ).

### Metformin improves the plasma endocrine profile of reproductive hormones

The supplementation of metformin in the diet significantly affected plasma progesterone concentrations when observed over time (Fig. 5A and 5B). From 30 weeks until 50 weeks of age, hens that received metformin in the diet tended to have lesser plasma progesterone concentrations when compared to the control. No difference was observed among treatment groups at 25 and 55 weeks of age. Interestingly, at 60 and 65 weeks of age, we observed a significant increase in plasma progesterone concentrations in hens that received metformin in the diet compared to the control, which is consistent with the prolonged egg production observed in these groups (Fig. 5A). When plasma progesterone concentrations were observed across time for each treatment group, we noted a significant decline in plasma progesterone concentrations in hens that received no metformin in the diet beginning at 50 weeks of age. However, plasma progesterone concentrations remained elevated and more consistent in hens supplemented with metformin in the diet at all levels (Fig. 5B). Like plasma progesterone concentrations, hens that received metformin in the diet had significantly lesser plasma total estrogen concentrations at 35, 40, 45, 50, and 60 weeks of age when compared to the control (Fig. 6A). When plasma total estrogen concentrations were studied over time, a similar trend was observed in which hens that received 50- or 75 mg/kg had a more consistent total estrogen profile over time when compared to the hens which received 0- or 25 mg/kg in the diet (Fig. 6B). Beginning at 30 weeks of age through 65 weeks of age, hens that received metformin in the diet had significantly lesser plasma testosterone concentrations when compared to the control (Fig. 7A). When plasma testosterone concentrations were studied over time, from 25 to 65 weeks of age, we observed consistent plasma testosterone concentration patterns within each treatment group (Fig. 7B). There was no effect of metformin on plasma androstenedione concentrations across all time points and over time (Fig. 8A and 8B). However, when we analyzed the testosterone to androstenedione ratio, a novel biomarker used to assess the metabolic phenotype in women with PCOS (Misichronis, G et al., 2011; Munzker, J et al., 2017; Dumesic, Daniel A et al., 2021), we observed that hens that received 50 or 75 mg/kg showed significantly lesser testosterone to androstenedione ratios at 40, 45, 50, 60 and 65 weeks of age when compared to the control (Fig. 9A). When the testosterone to androstenedione ratio is observed across time, hens that received metformin in the diet had a lower ratio that remained consistent compared to the control (Fig. 9B).

**Figure 6.**
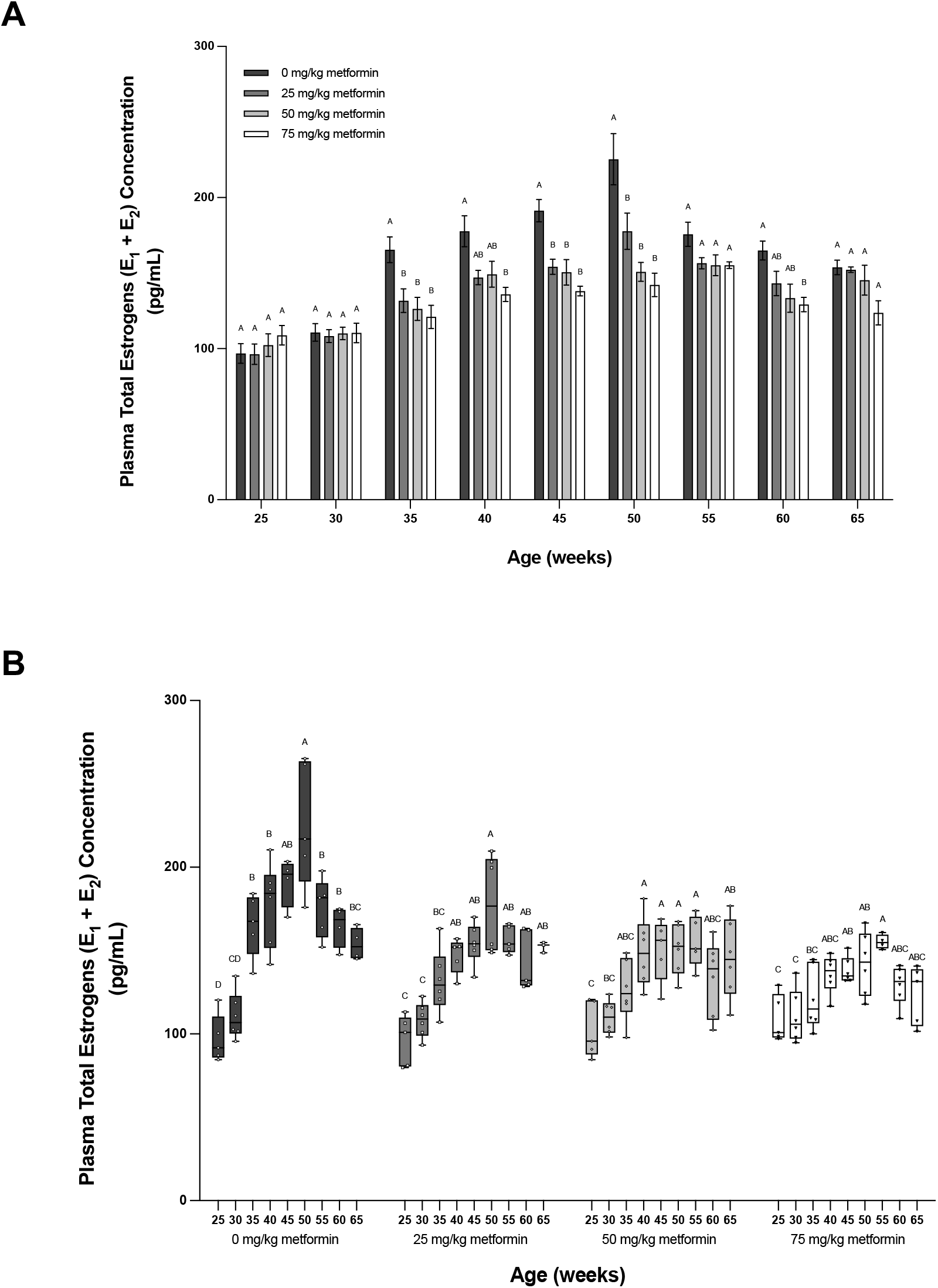
Effect of metformin supplementation on plasma total estrogen concentration. Broiler breeder hen diet was supplemented with metformin (0, 25, 50, or 75 mg/kg bw) from 25 to 65 weeks of age. Total estrogen concentrations (ng/mL) were determined by radioimmunoassay in blood samples collected from a subset of hens (n=6 hens/treatment group) every 5 weeks from 25-65 weeks of age **(A)**. Plasma total estrogen concentrations (ng/ml) within each treatment group over the course of the study is shown in **(B)**. Values are means ± SEM.^A, B,C, D^ P < 0.05 by one-way ANOVA; n= 6 hens/treatment group for each time point.

**Figure 7.**
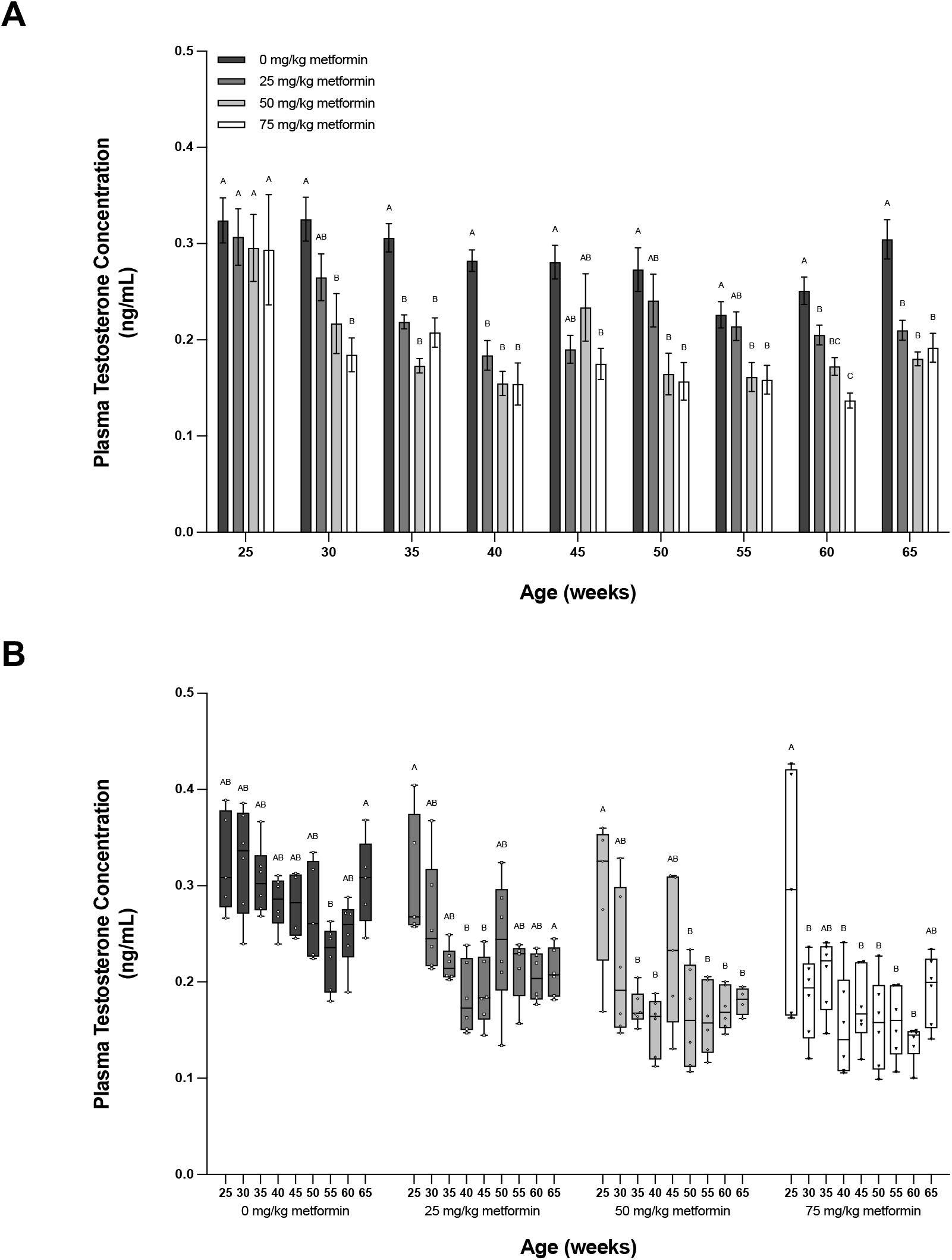
Effect of metformin supplementation on plasma testosterone concentration. Broiler breeder hen diet was supplemented with metformin (0, 25, 50, or 75 mg/kg bw) from 25 to 65 weeks of age. Testosterone concentrations (ng/mL) were determined by radioimmunoassay in blood samples collected from a subset of hens (n=6 hens/treatment group) every 5 weeks from 25-65 weeks of age **(A)**. Plasma testosterone concentrations (ng/ml) within each treatment group over the course of the study is shown in **(B).** Values are means ± SEM. ^A, B,^ P < 0.05 by one-way ANOVA; n= 6 hens/treatment group for each time point.

**Figure 8.**
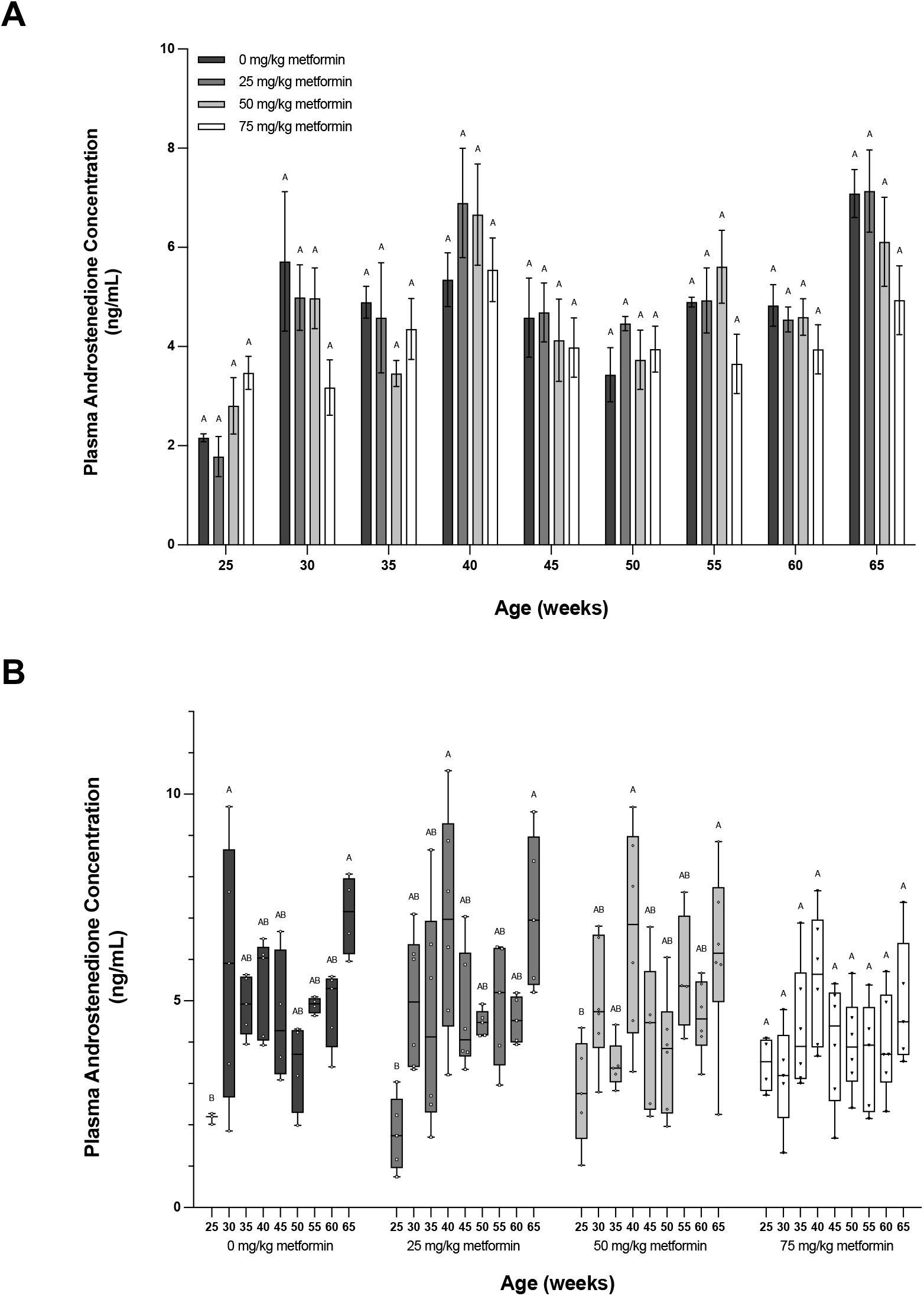
Effect of metformin supplementation on plasma androstenedione concentration. Broiler breeder hen diet was supplemented with metformin (0, 25, 50, or 75 mg/kg bw) from 25 to 65 weeks of age. Androstenedione concentrations (ng/mL) were determined by radioimmunoassay in blood samples collected from a subset of hens (n=6 hens/treatment group) every 5 weeks from 25-65 weeks of age **(A)**. Plasma androstenedione concentrations (ng/ml) within each treatment group over the course of the study is shown in **(B)**. Values are means ± SEM. ^A, B,^ P < 0.05 by one-way ANOVA; n= 6 hens/treatment group for each time point.

**Figure 9.**
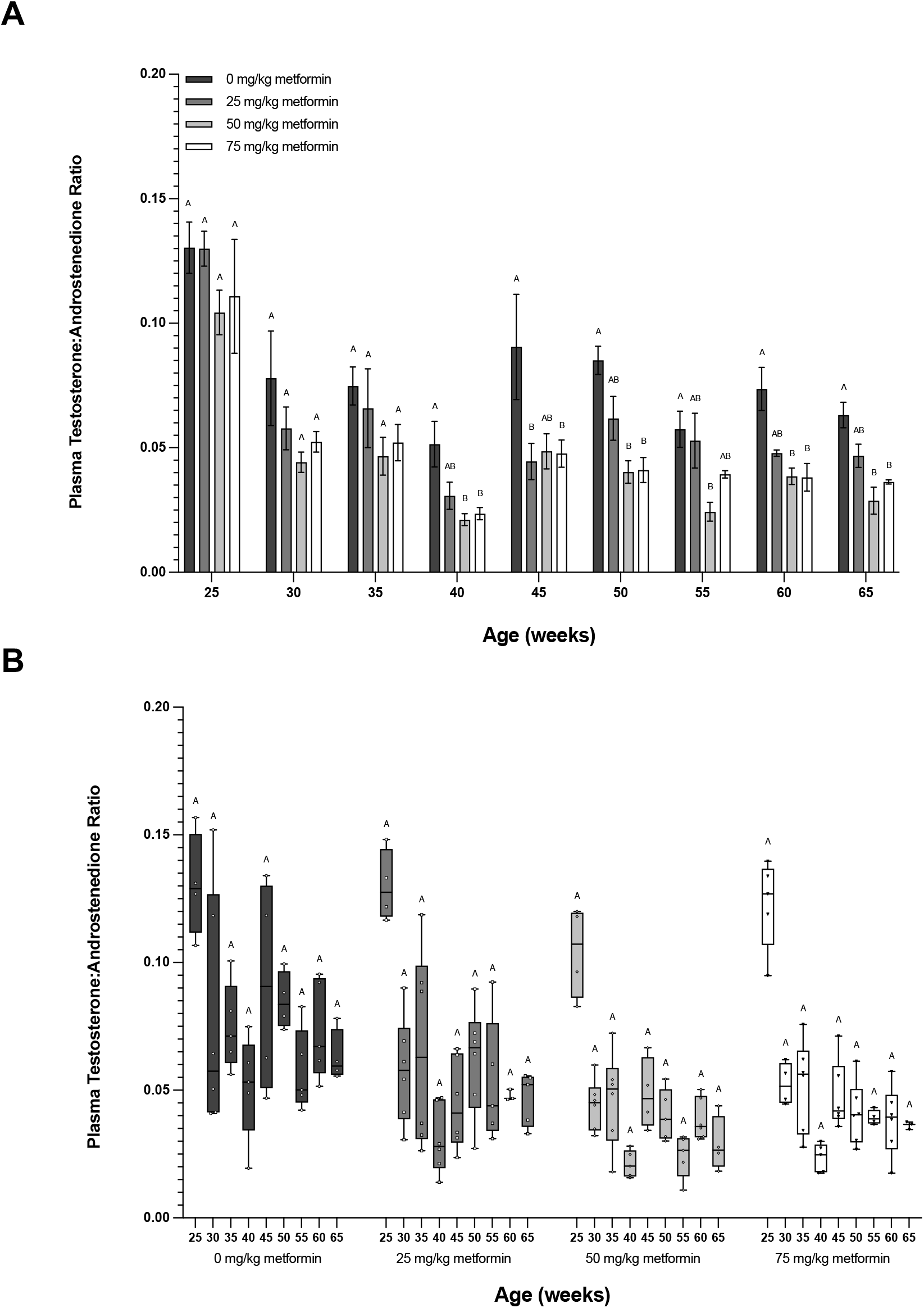
Effect of metformin supplementation on the plasma testosterone to androstenedione ratio. Broiler breeder hen diet was supplemented with metformin (0, 25, 50, or 75 mg/kg bw) from 25 to 65 weeks of age. The plasma testosterone to androstenedione ratio was calculated from the plasma androstenedione and testosterone concentrations (see Fig. 6 and 7) and represented 25-65 weeks of age **(A)** or within each treatment group over the course of the study **(B).** Values are means ± SEM. ^A, B^ P < 0.05 by one-way ANOVA; n= 6 hens/treatment group for each time point.

## DISCUSSION

We report for the first time that dietary metformin supplementation enhances ovarian function in the broiler breeder hen through the normalization of the ovarian follicular hierarchy and endocrine profile of reproductive hormones. The improved ovarian function was associated with decreased body weights, decreased accretion of abdominal adipose tissue, increased and prolonged egg production, maintained fertility and hatchability and altered plasma steroid hormone concentrations.

Our findings related to body weight are consistent with previous studies in which metformin was associated with a decrease in overall body weight of human subjects (Fleming et al., 2002; Bray et al., 2012; Tokubuchi et al., 2017; Guan et al., 2020; Chen et al., 2021), rodent (Jing et al., 2018; Rencbar et al., 2018; Coll et al., 2020) and Leghorn chicken (Chen et al., 2011). It has been previously reported that 40-80% of women diagnosed with PCOS are overweight or obese (Sam, 2007). In women with this condition, a decrease in body weight has been positively correlated with improved reproductive function, although some studies have suggested that weight loss may be the key to improving reproductive function rather than metformin itself (Tang et al., 2006a). Our findings also showed a significant decrease in abdominal adipose tissue weight in hens that received the highest dose of metformin. Consistent with our findings of a decreased fat pad weight, several studies mentioned above also showed that metformin leads to a decrease in the hip to waist circumference in women, which is consistent with a decrease in the accretion of visceral adipose tissue.

This is the first study to report that dietary metformin supplementation corrects the excessive recruitment of prehierarchical and preovulatory ovarian follicles, thus normalizing the ovarian follicular hierarchy in broiler breeder hens. Previous studies on the ovarian dysfunction encountered in broiler breeder hens have mostly focused on hens allowed to consume feed *ad libitum*, which is often associated with excessive follicular recruitment, increased incidence of double and internal ovulations, and premature ovarian regression. To correct these issues, the industry standard is to restrict feed intake to ‘improve’ ovarian function, but this feed restriction is often imprecise and hard to manage at the flock level. Although our hens were feed restricted according to industry standards, we still observed a greater number of hens exhibiting ovarian dysfunction in the group which received no metformin in the diet compared to hens which received metformin. The typical ovarian aberrations encountered in these hens were a disrupted follicular hierarchy (shown in Figure 2) and an increased incidence of internal ovulations. It is important to note that these anomalies were not present in all of the hens in this group, and some were able to maintain egg production and had healthy ovaries at the end of the study. While the aberrant recruitment of follicles and ovarian dysfunction encountered in the restricted-fed hens is similar to what is encountered in hens fed *ad libitum*, it is much less severe in restricted hens (typically only the hyper-recruitment of a few preovulatory follicles) compared to hens fed *ad libitum* (a complete double preovulatory hierarchy often encountered in these hens).

Interestingly, metformin supplementation, even at the lowest dose, was able to ameliorate these issues and ‘normalize’ the ovarian follicular hierarchy without observed effects on hen weight or abdominal fat pad weight in these hens. Although a decrease in body weight and the accretion of adipose tissue have been correlated with improved follicular hierarchies, previous studies have suggested that metformin can exert direct effects on the ovary (Ehrmann et al., 1997; Vrbikova et al., 2001; Bertoldo et al., 2014). The mechanisms underlying improved ovarian function in the broiler breeder hen still remain unclear so we cannot assume that loss of weight is a precondition to the improved follicular hierarchy observed in these hens. Future studies in the chicken should investigate the mechanisms in which metformin is able to improve the follicular hierarchy.

Consistent with our findings, previous studies have reported a ‘normalization’ of the ovarian follicular profile in both rats (Di Pietro et al., 2015; Rencbar et al., 2018; Ali et al., 2020) and humans (Fleming et al., 2002; Morley et al., 2017) and has been reported to increase ovulation rates (Nestler et al., 1998, 2002; Fleming et al., 2002) as well as a return to normal menstrual cyclicity (Velazquez et al., 1994; Morin-Papunen et al., 1998; Moghetti et al., 2000; van Santbrink et al., 2005) in women with PCOS.

Along with the ‘normalization’ of the ovarian follicular hierarchy, dietary metformin supplementation was also associated with increased and prolonged egg production, with the highest dose having the most significant effect. Even with strict feed restrictions, broiler breeder hens often have a significant reduction in egg production beginning around 45-50 weeks of age, and many cease laying by 60 weeks of age. We chose to begin metformin supplementation at 25 weeks of age as most hens should have begun to lay by this time. We observed a sharp decline in egg production in hens that received the highest dose of metformin around 27 weeks through 30 weeks of age (EA Weaver, R Ramachandran, unpublished data), which was not observed in a previous study conducted in our laboratory in which we supplemented metformin in the diet from 42-60 weeks of age (EA Weaver and R Ramachandran, unpublished data). Previous studies have shown that a certain percentage of body fat is required for hens to begin laying and then maintain egg production (Bornstein et al., 1983; Hurwitz and Plavnik, 1989; Yuan et al., 1994), so this observed decline in egg production could be due to the effect of metformin on body fat percentage in the hens which received the highest dose. Further studies are required to optimize the ideal body composition and age to begin supplementing metformin into the diet of broiler breeder hens. Although a significant decline in egg production was observed early on, these hens recovered egg production, which was maintained for the duration of the study. During the last 10 weeks of the study (55-65 weeks of age), we observed maintenance of egg production in hens that received metformin in their diet, even at the lowest dose of 25 mg/kg of body weight, with the two highest doses having the most significant effect. Interestingly, metformin supplementation appeared to push back the age of peak lay compared to hens which received no metformin. The age of peak lay was 27 weeks of age in hens which received no metformin in the diet, 33 weeks of age in hens which received 25 mg metformin/kg of body weight and 35 weeks of age for hens which received 50 or 75 mg metformin/kg body weight (data not shown). Overall, hens that received metformin laid a greater number of eggs on average which can be attributed to this maintenance of egg production. Based on previous studies, this maintenance of egg production could potentially be attributed to a reduction in body weight, reduction in the accumulation of abdominal adipose tissue, an effect of metformin on steroid hormone production or the direct effect of metformin on the ovary. However, further studies are necessary to determine the exact mechanisms by which metformin improves egg production in the broiler breeder hen.

We observed that metformin had no effect on fertility or hatchability amongst all treatment groups from 35 to 55 weeks of age. In hens which received no metformin in the diet or the lowest dose (25 mg metformin/kg body weight), a decline in both fertility and hatchability was observed beginning at 55 weeks of age through the end of the study which is expected based on current broiler breeder standards. Interestingly, the hens which received the two highest doses of metformin (50 or 75 mg/kg body weight) were able to maintain significantly higher rates of fertility and hatchability through the end of the study. Our findings are in agreement with previous studies which have shown that metformin can increase fertility and pregnancy rates (Velazquez et al., 1997; Nestler et al., 2002; Tang et al., 2006b; Morley et al., 2017; Tso et al., 2020) in women with PCOS. The mechanisms by which metformin is able to improve fertility and hatchability remain unknown and future studies should focus on the effects of metformin on egg quality and condition as these both play roles in hen fertility and hatchability.

Finally, we observed that supplementation of metformin in the diet significantly affected the plasma endocrine profile of reproductive hormones in broiler breeder hens. It has been previously reported that metformin can normalize the endocrine profile in both women (Velazquez et al., 1994; Xing et al., 2020) and rats (Zhang et al., 2017). In a previous study conducted in our laboratory, we reported that metformin attenuates steroidogenesis in the granulosa cells isolated from prehierarchical follicles and preovulatory follicles (Weaver and Ramachandran, 2020); however, this is the first report on the long-term *in vivo* effects of metformin in broiler breeder chickens. In the hen, the largest preovulatory follicle produces a large amount of progesterone, while prehierarchical follicles are major producers of estrogens.

We observed increased plasma progesterone and total estrogens concentrations in hens that received no metformin early on in the study, which is consistent with the hyper-recruitment of preovulatory and prehierarchical follicles. This increase in plasma total estrogens is not observed in hens which received metformin in the diet and concentrations remained consistent throughout the study. Interestingly, around 50 weeks of age, we observe a significant decline in plasma progesterone concentration and begin to see a decline in plasma total estrogen concentrations in hens that received no metformin in the diet. In contrast, progesterone concentrations remained consistent and elevated in hens that received metformin at all doses. As metformin supplementation maintained egg production during this time, this increase in plasma progesterone is consistent with maintaining a preovulatory follicular hierarchy. We observed a significant effect of metformin on plasma testosterone concentrations, consistent with previous studies that reported decreased plasma testosterone in women who received metformin (Nestler and Jakubowicz, 1996; Kurzthaler et al., 2014; Guan et al., 2020). Increased testosterone levels have been previously reported in broiler breeder hens allowed to consume feed *ad libitum* (Taherkhani et al., 2010; Singh et al., 2013); however, we are the first to report increased plasma testosterone even when restricted-fed. Based on our observations and similar to women with PCOS, it appears that an increased plasma testosterone profile is associated with increased body weight and adverse reproductive outcomes in the broiler breeder hen. In the present study, we found that plasma androstenedione concentrations in the broiler breeder hen treated with metformin did not differ. However, when comparing plasma testosterone to androstenedione ratios, we observed a significantly decreased testosterone to androstenedione ratio beginning at 40 weeks of age through the end of the study in hens that received metformin in the diet at 50 or 75 mg/kg of body weight. Previous studies have suggested that the androstenedione to testosterone ratio may be utilized as a biomarker of metabolic health in women with PCOS. An increased testosterone to androstenedione ratio has been associated with a higher likelihood of developing metabolic dysfunction, obesity, and insulin resistance in women with PCOS (Misichronis, G et al., 2011; Munzker, J et al., 2017; Dumesic, Daniel A et al., 2021).

In summary, we provide evidence that dietary metformin supplementation improves ovarian function in the broiler breeder hen. The improved ovarian function was associated with decreased body weights and abdominal fat pad weights, the normalization of the ovarian follicular hierarchy, increased and prolonged egg production, maintenance of fertility and hatchability and an improved endocrine profile of reproductive hormones. As the broiler breeder hen continues to be the poorest in reproductive efficiency of all avian species, a cheap and effective method to improve egg production is crucial to maintain broiler production and feed the growing population. Future studies are needed to determine the mechanisms that underlie the beneficial effect of metformin in correcting the ovarian follicular hierarchy and increasing egg production in the chicken.

## DECLARATION OF INTEREST

There is no conflict of interest that could be perceived as prejudicing the impartiality of the research reported.

## FUNDING

This project was supported by Agriculture and Food Research Initiative Competitive Grant no. 2017-67015-26506 from the USDA National Institute of Food and Agriculture and in part, by NIH Grant T32GM108563.

## AUTHOR CONTRIBUTION

EAW designed and conducted the trial and experiments and analyzed all data. EAW and RR wrote the manuscript.

## ACKNOWLEDGEMENTS

The authors wish to thank the staff at Poultry Education and Research Center at Penn State for their assistance with the trial.

## Notes

### Competing Interest Statement

The authors have declared no competing interest.

### Summary of Updates

Figures revised and new data added in Figure 4. The supplementary data has also replaced with the original Figure 1. Other revisions were made to the results and the discussion to include new data added in Figure 4. Other revisions made to the text were done to clarify methods and improve flow of discussion.

