## Supplementary Figure 1 for "Metformin Improves Ovarian Function and Increases Egg Production in Broiler Breeder Hens"

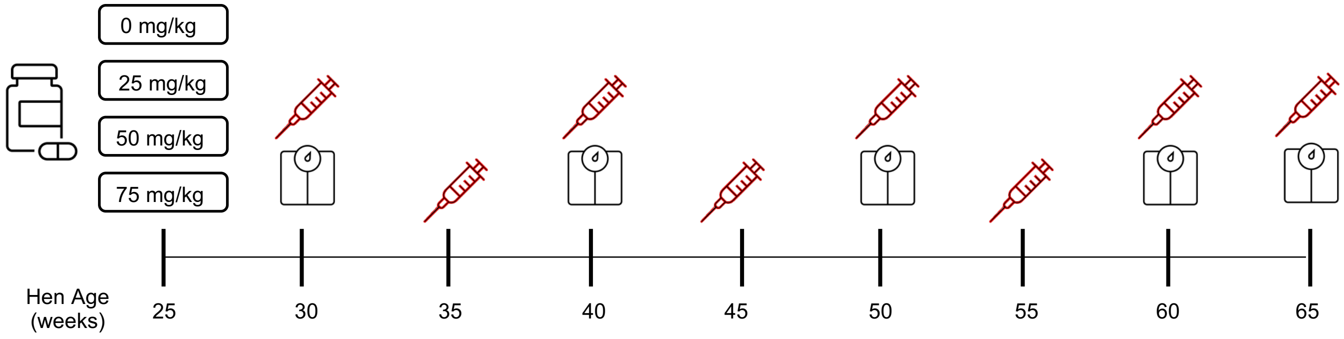


**Supplementary Figure 1. Experimental design and data collection timeline.** Broiler breeder hens were raised from day-old chicks and randomly assigned to a treatment group (n=45 hens/treatment group) to receive metformin (0, 25, 50 or 75 mg/kg body weight) in the diet starting from 25-65 weeks of age. Blood samples were collected from a subset of hens (n=6 hens/treatment group) every 5 weeks starting from 30 weeks of age. A subset of hens (n=10 hens/treatment group) were weighed every 10 weeks. At the end of the study, a subset of hens (n=12 hens/treatment group) were euthanized to collect tissues.
